# Scale matters: Large language models with billions (rather than millions) of parameters better match neural representations of natural language

**DOI:** 10.1101/2024.06.12.598513

**Authors:** Zhuoqiao Hong, Haocheng Wang, Zaid Zada, Harshvardhan Gazula, David Turner, Bobbi Aubrey, Leonard Niekerken, Werner Doyle, Sasha Devore, Patricia Dugan, Daniel Friedman, Orrin Devinsky, Adeen Flinker, Uri Hasson, Samuel A. Nastase, Ariel Goldstein

**Affiliations:** Department of Psychology and the Neuroscience Institute, Princeton University, Princeton, NJ; McGovern Institute for Brain Research, Massachusetts Institute of Technology, Cambridge, MA; New York University Grossman School of Medicine, New York, NY; Business School, Data Science Department and Cognitive Science Department, Hebrew University, Jerusalem, Israel

## Abstract

Recent research has used large language models (LLMs) to study the neural basis of naturalistic language processing in the human brain. LLMs have rapidly grown in complexity, leading to improved language processing capabilities. However, neuroscience researchers haven’t kept up with the quick progress in LLM development. Here, we utilized several families of transformer-based LLMs to investigate the relationship between model size and their ability to capture linguistic information in the human brain. Crucially, a subset of LLMs were trained on a fixed training set, enabling us to dissociate model size from architecture and training set size. We used electrocorticography (ECoG) to measure neural activity in epilepsy patients while they listened to a 30-minute naturalistic audio story. We fit electrode-wise encoding models using contextual embeddings extracted from each hidden layer of the LLMs to predict word-level neural signals. In line with prior work, we found that larger LLMs better capture the structure of natural language and better predict neural activity. We also found a logarithmic relationship where the encoding performance peaks in relatively earlier layers as model size increases. We also observed variations in the best-performing layer across different brain regions, corresponding to an organized language processing hierarchy.

## Introduction

How has the functional architecture of the human brain come to support everyday language processing? Modeling the underlying neural basis that supports natural language processing has proven to be prohibitively challenging for many years. Deep learning has brought about a transformative shift in our ability to model natural language in recent years. Leveraging principles from statistical learning theory and using vast real-world datasets, deep learning algorithms can reproduce complex natural behaviors in visual perception, speech analyses, and even human-like conversations. With the recent emergence of large language models (LLMs), we are finally beginning to see explicit computational models that respect and reproduce the context-rich complexity of natural language and communication. LLMs rely on simple self-supervised objectives (e.g., next-word prediction) to learn to produce context-specific linguistic outputs from real-world corpora—and, in the process, implicitly encode the statistical structure of natural language into a multidimensional embedding space (Linzen & Baroni, 2021; Manning et al., 2020; Pavlick, 2022).

Critically, there appears to be an alignment between the internal activity in LLMs for each word embedded in a natural text and the internal activity in the human brain while processing the same natural text. Indeed, recent studies have revealed that the internal, layer-by-layer representations learned by these models predict human brain activity during natural language processing better than any previous generations of models (Caucheteux & King, 2022; Goldstein et al., 2022, 2024; Kumar et al., 2024; Schrimpf et al., 2021).

LLMs, however, contain millions or billions of parameters, making them highly expressive learning algorithms. Combined with vast training text, these models can encode a rich array of linguistic structures—ranging from low-level morphological and syntactic operations to high-level contextual meaning—in a high-dimensional embedding space. Recent work has argued that the “size” of these models—the number of learnable parameters—is critical, as some linguistic competencies only emerge in larger models with more parameters (Bommasani et al., 2021; Kaplan et al., 2020; Manning et al., 2020; Sutton, 2019; C. Zhang et al., 2021). For instance, in-context learning (Liu et al., 2022; Xie et al., 2021) involves a model acquiring the ability to carry out a task for which it was not initially trained, based on a few-shot examples provided by the prompt. This capability is present in the bigger GPT-3 (Brown et al., 2020) but not in the smaller GPT-2, despite both models having similar architectures. This observation suggests that simply scaling up models produces more human-like language processing. Indeed, research in comparative neuroscience has suggested that uniquely human cognitive abilities emerged from scaling up the primate brain (Herculano-Houzel, 2012). While building and training LLMs with billions to trillions of parameters is an impressive engineering achievement, such artificial neural networks are tiny compared to cortical neural networks. In the human brain, each cubic millimeter of cortex contains a remarkable number of about 150 million synapses, and the language network can cover a few centimeters of the cortex (Cantlon & Piantadosi, 2024).

Our study focuses on one crucial question: What is the relationship between the size of an LLM and how well it can predict linguistic information encoded in the brain? In this study, we define “model size” as the number of all trainable parameters in the model. In addition to size, we also consider the model’s expressivity: its capacity to predict the statistics of natural language. Perplexity measures expressivity by evaluating the average level of surprise or uncertainty the model attributes to a sequence of words. Larger models possess a greater capacity for expressing linguistic structure, which tends to yield lower (better) perplexity scores (Radford et al., 2019). In this paper, we hypothesized that larger models that capture linguistic structure more accurately (lower perplexity) would better capture neural activity.

To test this hypothesis, we used electrocorticography (ECoG) to measure neural activity in ten epilepsy patient participants while they listened to a 30-minute audio podcast. Invasive ECoG recordings more directly measure neural activity than non-invasive neuroimaging modalities like fMRI, with much higher temporal resolution. We extracted contextual embeddings at each hidden layer from multiple families of transformer-based LLMs, including GPT-2, GPT-Neo, OPT, and Llama 2 (Black et al., 2022; Radford et al., 2019; Touvron et al., 2023; S. Zhang et al., 2022), and fit electrode-wise encoding models to predict neural activity for each word in the podcast stimulus. We found that larger language models, with greater expressivity and lower perplexity, better predicted neural activity (Antonello et al., 2023). This result was consistent across all model families. Critically, we then focus on a particular family of models (GPT-Neo), which span a broad range of sizes and are trained on the same text corpora. This allowed us to assess the effect of scaling on the match between LLMs and the human brain while keeping the size of the training set constant.

## Results

To investigate scaling effects between model size and the alignment of the model internal representations (embeddings) with brain activity, we utilized four families of transformer-based language models: GPT-2, GPT-Neo, OPT, and Llama 2 (Black et al., 2022; Radford et al., 2019; Touvron et al., 2023; S. Zhang et al., 2022). These models span 82 million to 70 billion parameters and 6 to 80 layers (Table 1). Different families of models vary in architectural details and are trained on different text corpora. To control for these confounding variables, we also focused on the GPT-Neo family (Gao et al., 2020) with a comprehensive range of models that vary only in size (but not training data), spanning 125 million to 20 billion parameters. For simplicity, we renamed the four models as “SMALL” (gpt-neo-125M), “MEDIUM” (gpt-neo-1.3B), “LARGE” (gpt-neo-2.7B), and “XL” (gpt-neox-20b).

**Table 1.** Summary of four families of open large language models: GPT-2, GPT-Neo, OPT, and Llama-2. Context length is the maximum context length for the model, ranging from 1024 to 4096 tokens. The model name is the model’s name as it appears in the transformers package from Hugging Face (Wolf et al., 2019). Model size is the total number of parameters; M represents million, and B represents billion. The number of layers is the depth of the model, and the hidden embedding size is the internal width.

| Model Family | Context Length | Model Name | Model Size | Number of Layers | Hidden Embedding Size |
| --- | --- | --- | --- | --- | --- |
| GPT2 | 1024 | distilgpt2 | 82 M | 6 | 768 |
|  |  | gpt2 | 124 M | 12 | 768 |
|  |  | gpt2-medium | 355 M | 24 | 1024 |
|  |  | gpt2-large | 774 M | 32 | 1280 |
|  |  | gpt2-xl | 1.5 B | 48 | 1600 |
| GPT-Neo | 2048 | EleutherAI/gpt-neo-125M | 125 M | 12 | 768 |
|  |  | EleutherAI/gpt-neo-1.3B | 1.3 B | 24 | 2048 |
|  |  | EleutherAI/gpt-neo-2.7B | 2.7 B | 32 | 2560 |
|  |  | EleutherAI/gpt-neox-20b | 20 B | 44 | 6144 |
| OPT | 2048 | facebook/opt-125m | 125 M | 12 | 768 |
|  |  | facebook/opt-350m | 350 M | 24 | 1024 |
|  |  | facebook/opt-1.3b | 1.3 B | 24 | 2048 |
|  |  | facebook/opt-2.7b | 2.7 B | 32 | 2560 |
|  |  | facebook/opt-6.7b | 6.7 B | 32 | 4096 |
|  |  | facebook/opt-13b | 13 B | 40 | 5120 |
|  |  | facebook/opt-30b | 30 B | 48 | 7168 |
|  |  | facebook/opt-66b | 66 B | 64 | 9216 |
| Llama-2 | 4096 | meta-llama/Llama-2-7b-hf | 7 B | 32 | 4096 |
|  |  | meta-llama/Llama-2-13b-hf | 13 B | 40 | 5120 |
|  |  | meta-llama/Llama-2-70b-hf | 70 B | 80 | 8192 |

We collected ECoG data from ten epilepsy patients while they listened to a 30-minute audio podcast (*So a Monkey and a Horse Walk into a Bar*, 2017). We extracted high-frequency broadband power (70–200 Hz) in 200 ms bins at lags ranging from -2000 ms to +2000 ms relative to the onset of each word in the podcast stimulus. We ran a preliminary encoding analysis using non-contextual language embeddings (GloVe; Pennington et al., 2014) to select a subset of 160 language-sensitive electrodes across the cortical language network of eight patients; all subsequent analyses were performed within this set of electrodes (Goldstein et al., 2022). Using a podcast transcription, we next extracted contextual embeddings from each hidden layer across the four families of autoregressive large language models. We used the maximum context length of each word for each language model. We constructed linear, electrode-wise encoding models using contextual embeddings from every layer of each language model to predict neural activity for each word in the stimulus. We estimated and evaluated the encoding models using a 10-fold cross-validation procedure: ridge regression was used to estimate a weight matrix for predicting word-by-word neural signals in 9 out of 10 contiguous training segments of the podcast; for each electrode, we then calculated the Pearson correlation between predicted and actual word-by-word neural signals for the left-out test segment of the podcast. We repeated this analysis for 161 lags from -2,000 ms to 2,000 ms in 25 ms increments relative to word onset (Fig. 1).

**Figure 1.**
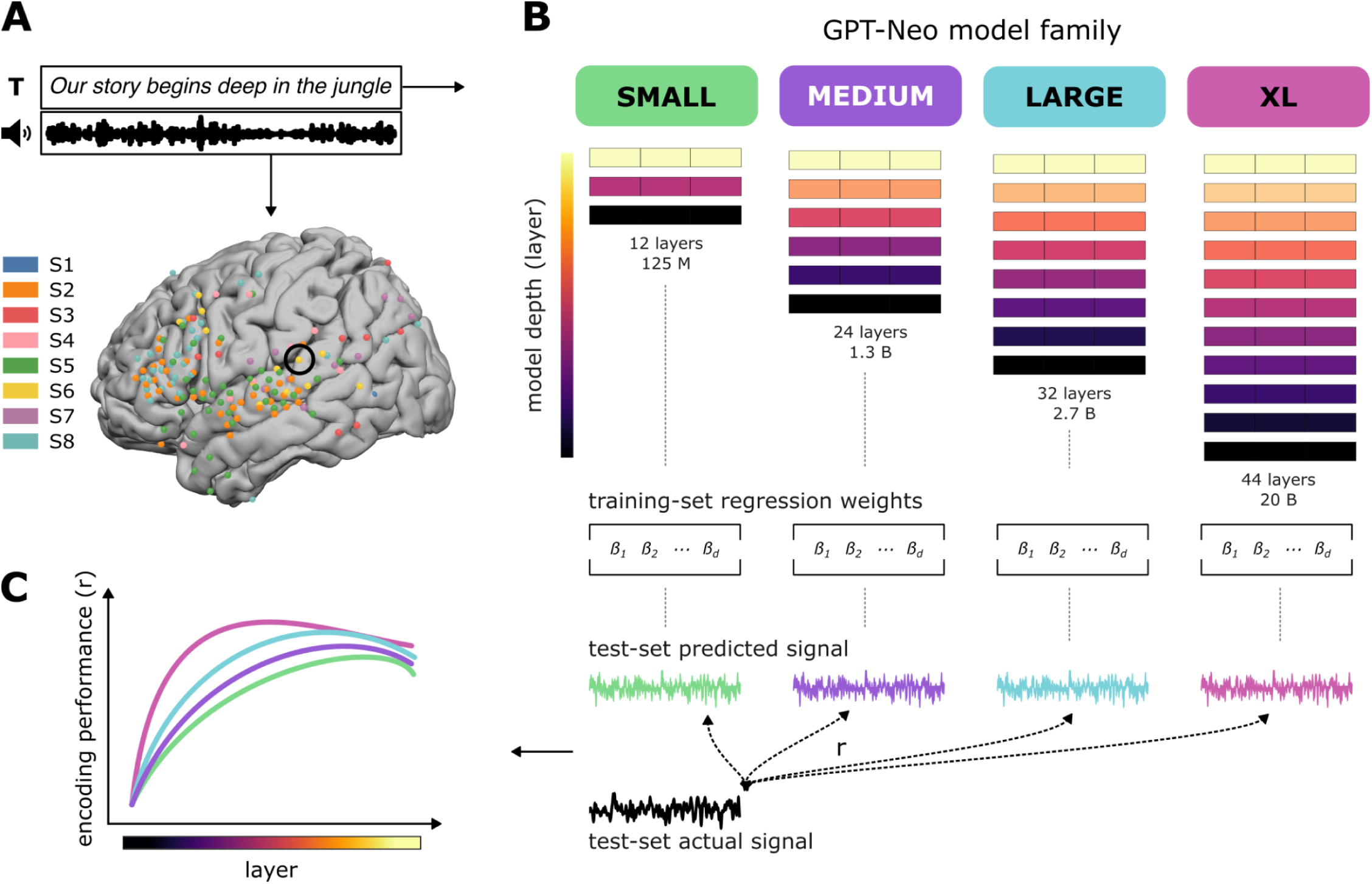
Naturalistic language comprehension model comparison framework. **A.** Participants listened to a 30-minute story while undergoing ECoG recording. A word-level aligned transcript was obtained and served as input to four language models of varying size from the same GPT-Neo family. **B.** For every layer of each model, a separate linear regression encoding model was fitted on a training portion of the story to obtain regression weights that can predict each electrode separately. Then, the encoding models were tested on a held-out portion of the story and evaluated by measuring the Pearson correlation of their predicted signal with the actual signal. **C.** Encoding model performance (correlations) was measured as the average over electrodes and compared between the different language models.

Prior to encoding analysis, we measured the “expressiveness” of different language models—that is, their capacity to predict the structure of natural language. Perplexity quantifies expressivity as the average level of surprise or uncertainty the model assigns to a sequence of words. A lower perplexity value indicates a better alignment with linguistic statistics and a higher accuracy during next-word prediction. For each model, we computed perplexity values for the podcast transcript. Consistent with prior research (Hosseini et al., 2022; Kaplan et al., 2020), we found that perplexity decreases as model size increases (Fig. 2A). In simpler terms, we confirmed that larger models better predict the structure of natural language.

**Figure 2.**
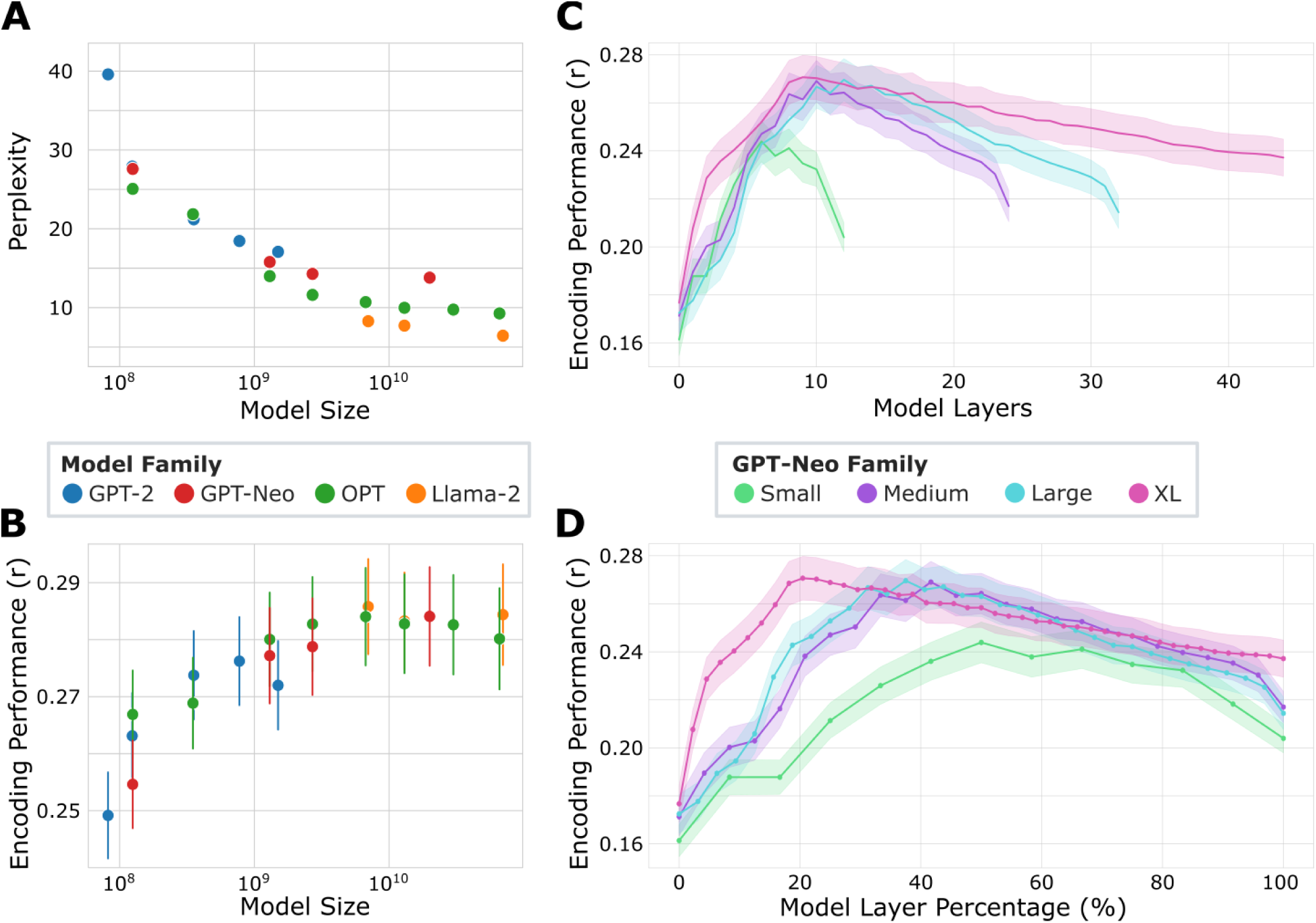
Model performance improves with increasing model size. **A.** The relationship between model size (measured as the number of parameters, shown on a log scale) and perplexity: as the model size increases, perplexity decreases. Each data point corresponds to a model. **B.** The relationship between model size (shown on a log scale) and brain encoding performance: correlations for each model are calculated by averaging the maximum correlations across all lags and layers across electrodes. As the model size increases, the encoding performance increases. Each data point corresponds to a model. The error bars represent standard error. **C.** For the GPT-Neo model family, the relationship between encoding performance and layer number. Encoding performance is best for intermediate layers. The shaded colors represent standard error. **D**. Same as C, but the layer number was transformed to a layer percentage for better model comparison.

### Larger language models better predict brain activity

We compared encoding model performance across language models at different sizes. For each electrode, we obtained the maximum encoding performance correlation across all lags and layers, then averaged these correlations across electrodes to derive the overall maximum correlation for each model (Fig. 2B). Using ECoG neural signals with superior temporal resolution, we replicated the previous fMRI work reporting a logarithmic relationship between model size and encoding performance (Antonello et al., 2023), indicating that larger models better predict neural activity. We also observed a plateau in the maximal encoding performance, occurring around 7 billion parameters (Fig. 2B), with a decline in performance for the OPT-66B model (Fig. S1). The size of the contextual embedding varies across models depending on the model’s size and architecture. This can range from 762 in the smallest distill GPT2 model to 8192 in the largest LLAMA-2 70 billion parameter model. To control for the different embedding dimensionality across models, we standardized all embeddings to the same size using principal component analysis (PCA) and trained linear encoding models using ordinary least-squares (OLS) regression, replicating the logarithmic relationship but with significantly lower encoding performance overall (Fig. S2). The PC features are used by the OLS models only.

To dissociate model size and control for other confounding variables, we next focused on the GPT-Neo models and assessed layer-by-layer and lag-by-lag encoding performance. For each layer of each model, we identified the maximum encoding performance correlation across all lags and averaged this maximum correlation across electrodes (Fig. 2C). Additionally, we converted the absolute layer number into a percentage of the total number of layers to compare across models (Fig. 2D). We found that correlations for all four models typically peak at intermediate layers, forming an inverted U-shaped curve, corroborating with previous fMRI findings (Caucheteux et al., 2021; Schrimpf et al., 2021; Toneva & Wehbe, 2019). Furthermore, we replicated the phenomenon observed by (Antonello et al., 2023), wherein smaller models (e.g. SMALL) achieve maximum encoding performance approximately three-quarters into the model, while larger models (e.g. XL) peak in relatively earlier layers before gradually declining. Leveraging the high temporal resolution of ECoG, we compared the encoding performance of models across various lags relative to word onset. We identified the optimal layer for each electrode and model and then averaged the encoding performance across electrodes. We found that XL significantly outperformed SMALL in encoding models for most lags from 2000 ms before word onset to 575 ms after word onset (Fig. S3). To establish a general baseline for encoding performance, we built encoding models using embeddings from the SMALL model with randomly initialized weights. The trained SMALL model exhibits significantly higher encoding performance across all layers compared to the untrained SMALL model (Fig. S4). We also assessed the encoding performance of contextual embeddings from LLMs against classic speech features and static GloVe embeddings (Table S1). The SMALL and XL embeddings achieved markedly higher encoding correlations than the speech features and GloVe embeddings (Fig. S5). We also built encoding models using subsets of the data and found that encoding performance increases as the volume of training data increases (Fig. S6).

### Encoding model performance across electrodes and brain regions

Next, we examined the differences in the encoding model across electrodes and brain regions. For each of the 160 electrodes, we identified the maximum encoding performance correlation across all lags and layers (Fig. 3A). Consistent with prior studies (Goldstein et al., 2022, 2025), our encoding model for SMALL achieved the highest correlations in superior temporal gyrus (STG) and inferior frontal gyrus (IFG). We then compared the encoding performances between SMALL and the other three models, plotting the percent change in encoding performance relative to SMALL for each electrode (Fig. 3B). Across all three comparisons, we observed significantly higher encoding performance for the larger models in approximately one-third of the 160 electrodes (two-sided pairwise t-test across cross-validation folds for each electrode, *p* < 0.05, Bonferroni corrected).

**Figure 3.**
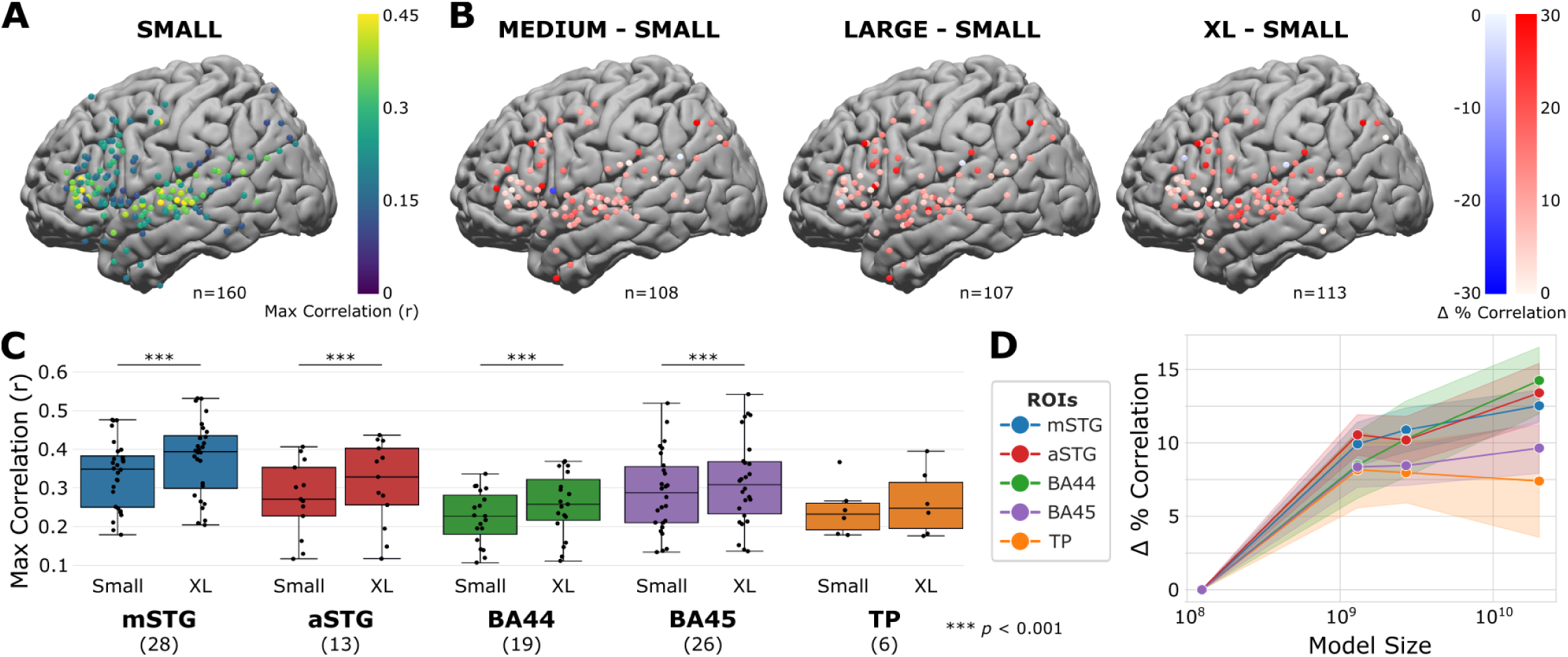
Model performance improves with increasing model size across electrodes and ROIs. **A.** Maximum correlation per electrode for SMALL. The encoding model achieves the highest correlations in STG and IFG. **B.** For MEDIUM, LARGE, and XL, the percentage difference in correlation relative to SMALL for all electrodes with significant encoding differences. The encoding performance is significantly higher for the bigger models for almost all electrodes across the brain (pairwise t-test across cross-validation folds). **C.** Maximum encoding correlations for SMALL and XL for each ROI (mSTG, aSTG, BA44, BA45, and TP area). The encoding performance is significantly higher for XL for all ROIs except TP. Each data point corresponds to an electrode in the corresponding ROI. **D.** Percent difference in correlation relative to SMALL for all ROIs. As model size increases, the percent change in encoding performance also increases for mSTG, aSTG, and BA44. After the medium model, the percent change in encoding performance plateaus for BA45 and TP. The shaded colors represent standard error.

We then compared the maximum correlations between SMALL and XL models across five regions of interest (ROIs) across the cortical language network (Fig. S7): middle superior temporal gyrus (mSTG, n = 28 electrodes), anterior superior temporal gyrus (aSTG, n = 13 electrodes), Brodmann area 44 (BA44, n = 19 electrodes), Brodmann area 45 (BA45, n = 26 electrodes), and temporal pole (TP, n = 6 electrodes). Encoding performance for the XL model significantly surpassed that of the SMALL model in mSTG, aSTG, BA44, and BA45 (Fig. 3C, Table S2). Additionally, we calculated the percent change in encoding performance relative to SMALL for each brain region by averaging across electrodes and plotting against model size (Fig. 3D). As model size increases, the fit to the brain nominally increases across all observed regions. However, the increase plateaued after the Medium model for regions BA45 and TP.

### The best layer for encoding performance varies with model size

In the previous analyses, we observed that encoding performance peaks at intermediate to later layers for some models and relatively earlier layers for others (Fig. 1C, 1D). To examine this phenomenon more closely, we selected the best layer for each electrode based on its maximum encoding performance across lags. To account for the variation in depth across models, we computed the best layer as the percentage of each model’s overall depth. We found that as models increase in size, peak encoding performance tends to occur in relatively earlier layers, being closer to the input in larger models (Fig. 4A). This was consistent across multiple model families, where we found a logarithmic relationship between model size and best encoding layers (Fig. 4B).

**Figure 4.**
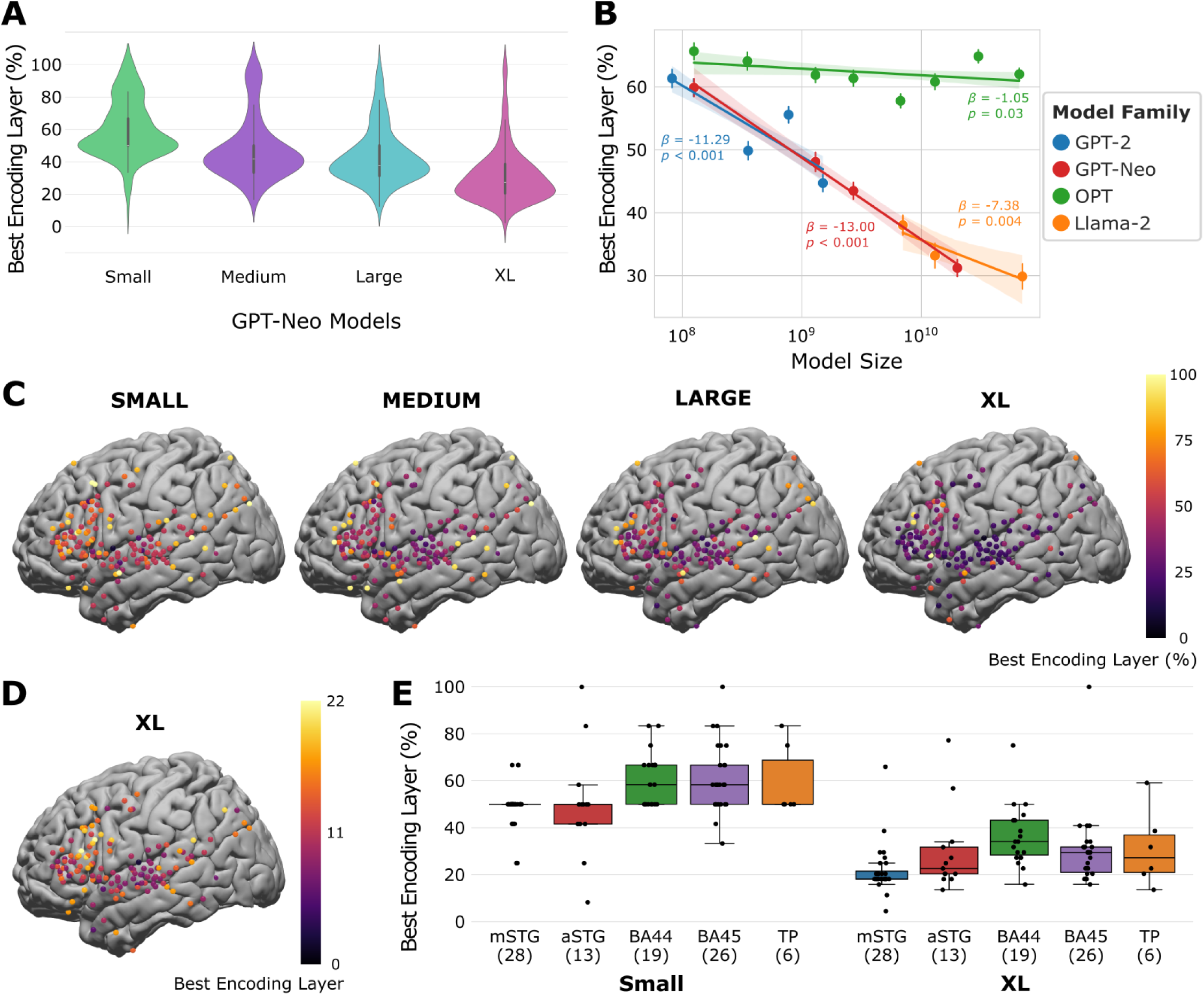
Relative layer preference varies with model size. **A.** Relative layer (in percentage of total number of layers) with peak encoding performance for all four GPT-Neo models: the larger the model size, the earlier relative layer where the encoding performance peaks. **B.** The relationship between model size (shown on a log scale) and best encoding layer (in percentage) for all four model families: as the model size increases, the best encoding layer (in percentage) decreases, although the rate of decrease is different between model families. We estimate a linear regression model per model family of the form: best percent layer ∼ log(model size). The slopes (*β*) indicate the decrease in the relative best-performing layer at increasing log model size; p-values are obtained from a Wald test against the null hypothesis that the slope is 0. Each data point corresponds to a model. **C.** Best relative encoding layer (in percentage) for all four GPT-Neo models. **D**. Best encoding layer for XL with electrodes that peak in the first half of the model (Layer 0 to 22). **E.** Best encoding layer (in percentage) for SMALL and XL for each ROI (mSTG, aSTG, BA44, BA45, and TP). Each data point corresponds to an electrode in the corresponding ROI.

We further observed variations of the best encoding layers across the brain within the same model. We found that the language processing hierarchy was better reflected in the best encoding layer preference for smaller than for larger models (Fig. 4C). Specifically, in the SMALL model, peak encoding was observed in earlier layers for STG electrodes and in later layers for IFG electrodes (Fig.4C, Table S3). A similar trend is evident in MEDIUM and partially in LARGE models, but not in the XL model, where the majority of electrodes exhibited peak encoding in the first 25% of all layers (Fig. 4C). However, despite the XL model showing less variance in the best layer distributions across cortex, we found the same hierarchy present for the first half of the model (layers 0–22, Fig. 4D). In this analysis, we observed that the best relative layer nominally increases from mSTG electrodes (M = 21.916, SD = 10.556) to aSTG electrodes (M = 29.720, SD = 17.979) to BA45 (M = 30.157, SD = 16.039) and TP electrodes (M = 31.061, SD = 16.305), and finally to BA44 electrodes (M = 36.962, SD = 13.140, Fig. 4E).

### The best lag for encoding performance does not vary with model size

Leveraging the high temporal resolution of ECoG, we investigated whether peak lag for encoding performance relative to word onset is affected by model size. For each ROI, we identified the optimal layer for each electrode in the ROI and then averaged the encoding performance (Fig. 5A). Within the SMALL model, we observed a trend where putatively lower-level regions of the language processing hierarchy peak earlier relative to word onset: mSTG encoding performance peaks around 25 ms before word onset, followed by aSTG encoding peak 225 ms after onset, and subsequently TP, BA44, and BA45 peak at approximately 350 ms. Within the XL model, we observed a similar trend, with mSTG encoding peaking first, followed by aSTG encoding peak, and finally, TP, BA44, and BA45 encodings. We also identified the lags when the encoding performance peaks for each electrode and visualized them on the brain map (Fig. 5B). Notably, the optimal lags for each electrode do not exhibit significant variation when transitioning from SMALL to XL.

**Figure 5.**
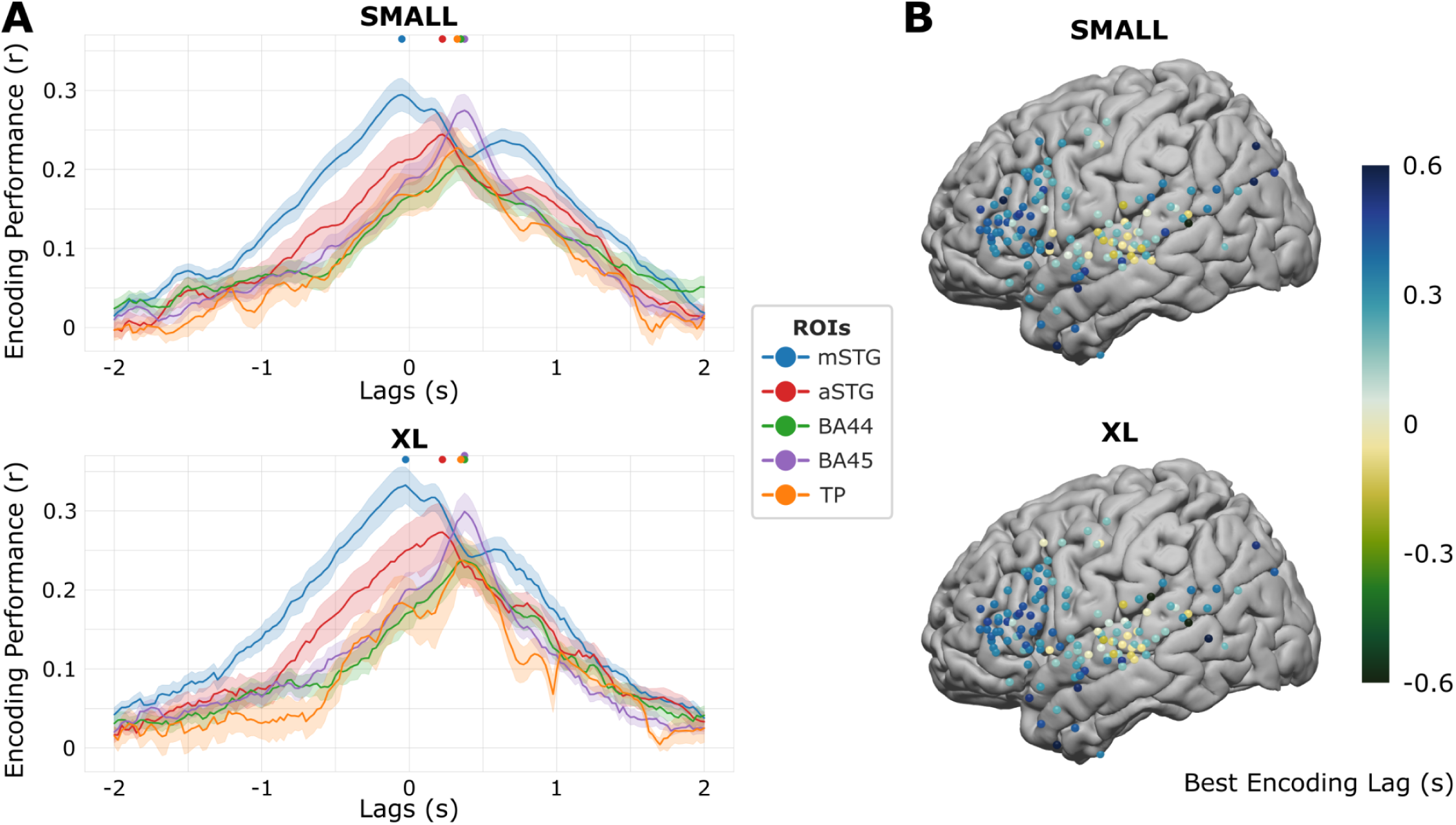
Encoding performance across lags does not vary with model size. **A.** Average ROI encoding performance for SMALL and XL models. mSTG encoding peaks first before word onset, then aSTG peaks after word onset, followed by BA44, BA45, and TP encoding peaks at around 400 ms after onset. The dots represent the peak lag for each ROI. **B.** Lag with best encoding performance correlation for each electrode, using SMALL and XL model embeddings. Only electrodes with the best lags that fall within 600 ms before and after word onset are plotted.

## Discussion

In this study, we investigated how the quality of model-based predictions of neural activity scales with LLM model size (i.e., the number of parameters across layers). Prior studies have shown that encoding models constructed from the internal embeddings of LLMs provide remarkably good predictions of neural activity during natural language comprehension (Caucheteux & King, 2022; Goldstein et al., 2022; Schrimpf et al., 2021). Corroborating prior work using fMRI (Antonello et al., 2023), across a range of models with 82 million to 70 billion parameters, we found that larger models are better aligned with neural activity. This result was consistent across several autoregressive LLM families varying in architectural details and training corpora and within a single model family trained on the same corpora and varying only in size. We suspect that the improved alignment with brain activity in larger models is driven by their increased expressivity and sensitivity to nuanced linguistic structure present in large-scale naturalistic datasets (Antonello et al., 2023). Our findings indicate that this trend does not trivially result from arbitrarily increasing model complexity: (a) models of varying size were estimated using regularized regression and evaluated using out-of-sample prediction to minimize the risks of overfitting, and (b) we obtained qualitatively similar results with explicitly matched dimensionality using PCA. Combined with the observation that larger models yield lower perplexity, our findings suggest that larger models’ capacity for learning the structure of natural language also yields better predictions of brain activity. By leveraging their increased size and representational power, these models have the potential to provide valuable insights into the mechanisms underlying language comprehension.

We focused on a particular family of models (GPT-Neo) trained on the same corpora and varying only in size to investigate how model size impacts layerwise encoding performance across lags and ROIs. We found that model-brain alignment improves consistently with increasing model size across the cortical language network. However, the increase plateaued after the MEDIUM model for regions BA45 and TP, possibly due to already high encoding correlations for the SMALL model and a small number of electrodes in the area, respectively.

A more detailed investigation of layerwise encoding performance revealed a logarithmic relationship where peak encoding performance tends to occur in relatively earlier layers as both model size and expressivity increase (Mischler et al., 2024). This is an unexpected extension of prior work on both language (Caucheteux & King, 2022; Kumar et al., 2024; Schrimpf et al., 2021; Toneva & Wehbe, 2019) and vision (Jiahui et al., 2023), where peak encoding performance was found at late-intermediate layers. Moreover, we observed variations in best relative layers across different brain regions, corresponding to a language processing hierarchy. This is particularly evident in smaller models and early layers of larger models. The inverted U-shaped trend of encoding performance commonly found in previous research is likely due to a “two-phase abstraction process” within LLMs (Cheng & Antonello, 2024; Csordás et al., 2025). In the early and intermediate layers of the model, a composition phase occurs, where low-level input features become increasingly abstract and contextualized. The intermediate layers of the model show the highest correlation with brain activity, presumably because they capture complex semantic and contextual information in a way that generalizes well across a variety of tasks (including prediction of human neural activity) (Antonello & Huth, 2024). Subsequently, a prediction phase happens in the later layers of the model, where the representations become more specialized for the LLM’s specific training objective (e.g., next-word prediction). This specialization can effectively constrict the more generalized feature representations, making these layers less optimal for predicting brain activity. Our results indicate that the initial composition phase does not take up more layers as models scale up in size. Larger models develop the necessary rich, abstract representations in the same number of layers as smaller models. Thus, as LLMs increase in size, the later layers of the model may contain representations that are increasingly divergent from the more general linguistic processing captured in brain activity. It is also possible that the later layers of larger models are overall underutilized and may not significantly contribute to benchmark performances during inference (Csordás et al., 2025; Fan et al., 2024; Gromov et al., 2024). Leveraging the high temporal resolution of ECoG, we found that putatively lower-level regions of the language processing hierarchy peak earlier than higher-level regions. However, we did not observe variations in the optimal lags for encoding performance across different model sizes. Since we exclusively employ textual LLMs, which lack inherent temporal information due to their discrete token-based nature, future studies utilizing multimodal LLMs integrating continuous audio or video streams, like Whisper or WavLM, may better unravel the relationship between model size and temporal dynamic representations in LLMs (Goldstein et al., 2025; Millet et al., 2023; Vaidya et al., 2022).

Our podcast stimulus comprised ∼5,000 words over a roughly 30-minute episode. Although this is a rich language stimulus, naturalistic stimuli of this kind have relatively low power for modeling infrequent linguistic structures (Hamilton & Huth, 2020). While perplexity for the podcast stimulus continued to decrease for larger models, we observed a plateau in predicting brain activity for the largest LLMs. The largest models learn to capture relatively nuanced or rare linguistic structures, but these may occur too infrequently in our stimulus to capture much variance in brain activity. Using a subsampling approach, we found that encoding performance also scales with the volume (and diversity) of language stimuli. Encoding performance may continue to increase for the largest models with more extensive stimuli (Antonello et al., 2023), motivating future work to pursue dense sampling with numerous, diverse naturalistic stimuli (Goldstein et al., 2025; LeBel et al., 2023).

The advent of deep learning has marked a tectonic shift in how we model brain activity in more naturalistic contexts, such as real-world language comprehension (Hasson et al., 2020; Richards et al., 2019). Traditionally, neuroscience has sought to extract a limited set of interpretable rules to explain brain function. However, deep learning introduces a new class of highly parameterized models that can challenge and enhance our understanding. The vast number of parameters in these models allows them to achieve human-like performance on complex tasks like language comprehension and production. It is important to note that LLMs have fewer parameters than the number of synapses in any human cortical functional network. Furthermore, the complexity of what these models learn enables them to process natural language in real-life contexts as effectively as the human brain does. Thus, the explanatory power of these models is in achieving such expressivity based on relatively simple computations in pursuit of a relatively simple objective function (e.g., next-word prediction). As in the human brain, while scaling alone may yield emergent cognitive abilities (Cantlon & Piantadosi, 2024; Herculano-Houzel, 2012), specialized architectural features likely also play a critical role (Friederici & Becker, 2025). As we continue to develop larger, more sophisticated models, the scientific community is tasked with advancing a framework for understanding these models to better understand the intricacies of the neural code that supports natural language processing in the human brain.

## Materials and Methods

### Participants

Ten patients (6 female, 20-48 years old) with treatment-resistant epilepsy undergoing intracranial monitoring with subdural grid and strip electrodes for clinical purposes participated in the study. Two patients consented to have an FDA-approved hybrid clinical research grid implanted, which includes standard clinical electrodes and additional electrodes between clinical contacts. The hybrid grid provides a broader spatial coverage while maintaining the same clinical acquisition or grid placement. All participants provided informed consent following the protocols approved by the Institutional Review Board of the New York University Grossman School of Medicine. The patients were explicitly informed that their participation in the study was unrelated to their clinical care and that they had the right to withdraw from the study at any time without affecting their medical treatment. One patient was removed from further analyses due to excessive epileptic activity and low SNR across all experimental data collected during the day.

### Stimuli

Participants listened to a 30-minute auditory story stimulus, “So a Monkey and a Horse Walk Into a Bar: Act One, Monkey in the Middle,” (*So a Monkey and a Horse Walk into a Bar*, 2017) from the This American Life Podcast. The audio narrative is 30 minutes long and consists of approximately 5000 words. Participants were not explicitly aware that we would examine word prediction in our subsequent analyses. The onset of each word was marked using the Penn Phonetics Lab Forced Aligner (Yuan & Liberman, 2008) and manually validated and adjusted as needed. The stimulus and alignment processes are described in prior work (Goldstein et al., 2022). In this study, we use the term “structure” to refer to a variety of linguistic patterns (e.g., morphology, syntax, semantics, context) that LLMs encode.

### Data acquisition and preprocessing

Across all patients, 1106 electrodes were placed on the left and 233 on the right hemispheres (signal sampled at or downsampled to 512 Hz). Brain activity was recorded from a total of 1339 intracranially implanted subdural platinum-iridium electrodes embedded in silastic sheets (2.3mm diameter contacts, Ad-Tech Medical Instrument; for the hybrid grids, 64 standard contacts had a diameter of 2 mm and an additional 64 contacts were 1mm diameter, PMT corporation, Chananssen, MN). We also preprocessed the neural data to get the power in the high-gamma-band activity (70-200 HZ). The full description of ECoG recording procedure is provided in prior work (Goldstein et al., 2022).

Electrode-wise preprocessing consisted of four main stages: First, large spikes exceeding four quartiles above and below the median were removed, and replacement samples were imputed using cubic interpolation. Second, the data were re-referenced using common average referencing. Third, 6-cycle wavelet decomposition was used to compute the high-frequency broadband (HFBB) power in the 70–200 Hz band, excluding 60, 120, and 180 Hz line noise. In addition, the HFBB time series of each electrode was log-transformed and z-scored. Fourth, the signal was smoothed using a Hamming window with a kernel size of 50 ms. The filter was applied in both the forward and reverse directions to maintain the temporal structure. Additional preprocessing details can be found in prior work (Goldstein et al., 2022).

### Electrode selection

We used a nonparametric statistical procedure with correction for multiple comparisons(Nichols & Holmes, 2002) to identify significant electrodes. We randomized each electrode’s signal phase at each iteration by sampling from a uniform distribution. This disconnected the relationship between the words and the brain signal while preserving the autocorrelation in the signal. We then performed the encoding procedure for each electrode (for all lags). We used GloVe embeddings for electrode selection to avoid biasing our main results toward a particular LLM. We repeated the encoding process 5000 times. After each iteration, the encoding model’s maximal value across all lags was retained for each electrode. We then took the maximum value for each permutation across electrodes. This resulted in a distribution of 5000 values, which was used to determine the significance for all electrodes. For each electrode, a *p*-value was computed as the percentile of the non-permuted encoding model’s maximum value across all lags from the null distribution of 5000 maximum values. Performing a significance test using this randomization procedure evaluates the null hypothesis that there is no systematic relationship between the brain signal and the corresponding word embedding. This procedure yielded a *p*-value per electrode, corrected for the number of models tested across all lags within an electrode. To further correct for multiple comparisons across all electrodes, we used a false-discovery rate (FDR). Electrodes with *q*-values less than .01 are considered significant. This procedure identified 160 electrodes from eight patients in the left hemisphere’s early auditory, motor cortex, and language areas.

### Perplexity

We computed the perplexity values for each LLM using our story stimulus, employing a stride length half the maximum token length of each model (stride 512 for GPT-2 models, stride 1024 for GPT-Neo models, stride 1024 for OPT models, and stride 2048 for Llama-2 models). These stride values yield the lowest perplexity value for each model. We also replicated our results on fixed stride length across model families (stride 512, 1024, 2048, 4096).

### Contextual embeddings

We extracted contextual embeddings from all layers of four families of autoregressive large language models. The GPT-2 family, particularly *gpt2-xl*, has been extensively used in previous encoding studies (Goldstein et al., 2022; Schrimpf et al., 2021). Here we include distilGPT-2 as part of the GPT-2 family. The GPT-Neo family, released by EleutherAI (Black et al., 2022), features three models plus GPT-Neox-20b, all trained on the Pile dataset (Gao et al., 2020). The OPT and Llama-2 families are released by MetaAI (Touvron et al., 2023; S. Zhang et al., 2022). For Llama-2, we use the pre-trained versions before any reinforcement learning from human feedback. All models we used are implemented in the HuggingFace environment (Wolf et al., 2019). We define “model size” as the combined width of a model’s hidden layers and its number of layers, determining the total parameters. We first converted the words from the raw transcript (including punctuation and capitalization) to tokens comprising whole words or sub-words (e.g., (1) there’s → (1) there (2) ’s). All models within the same model family adhere to the same tokenizer convention, except for GPT-Neox-20B, which utilizes a different tokenizer (Black et al., 2022). For each token, we utilized a context window with the maximum context length of each language model containing prior tokens from the podcast (i.e., the token and its history) and extracted the embedding for the final token in the sequence (i.e., the token itself). To facilitate a fair comparison of the encoding effect across different models, we aligned all tokens in the story across all models. We averaged the token embeddings if a word is split into several tokens, resulting in one embedding per word for each model.

Transformer-based language model consists of blocks containing a self-attention sub-block and a subsequent feedforward sub-block. The output of a block is obtained through a residual connection applied to the sum of the block’s input and the output of the feedforward sub-block. The self-attention output is added to this sum later in the layer normalization step. This output is commonly referred to as the “hidden state” of language models. This hidden state is considered the contextual embedding for the preceding block. For convenience, we refer to the blocks as “layers”; that is, the hidden state at the output of block 3 is referred to as the contextual embedding for layer 3. To generate the contextual embeddings for each layer, we store each layer’s hidden state for each word in the input text. Fortunately, the HuggingFace implementation of those language models automatically stores these hidden states when a forward pass of the model is conducted. Different models have different numbers of layers and embeddings of different dimensionality. For instance, gpt-neo-125M (SMALL) has 12 layers, and the embeddings at each layer are 768-dimensional vectors. Since we generate an embedding for each word at every layer, this results in 12 768-dimensional embeddings per word.

### Static embeddings

We constructed static embeddings with classic speech features and GloVe to compare with the encoding performance of LLMs. First, we extracted features capturing lower-level acoustic qualities of speech. Using the stimulus transcript as input, we created one-hot vectors for phonetic and articulatory features. Phoneme classes (39 total classes) were obtained from the Carnegie Mellon Pronouncing Dictionary (*The CMU Pronouncing Dictionary*, n.d.). We further classified the phonemes based on their place of articulation (9 classes), manner of articulation (9 classes), and voiced or voiceless status (3 classes), according to the general American English consonants of the International Phonetic Alphabet. Given that each word consists of multiple phonemes, we averaged the one-hot vectors for all phonetic and articulatory features for each word. Second, we extracted linguistic features using spaCy (Honnibal et al., 2020), including part of speech (17 classes), tag (50 classes), function or content word (3 classes), dependency (45 classes), whether the word is an alpha character (binary), and whether the word is a stop word (binary). We also extracted prefix (30 classes) and suffix (44 classes) information using the Cambridge Dictionary. We constructed one-hot vectors for each multi-class feature and one-dimensional vectors for each binary feature. Third, for each word, we obtained word frequency from the Google Web Trillion Word Corpus (Brants & Franz, 2006) and from our own dataset. Fourth, we generated static word embeddings of dimension 50 using GloVe (Pennington et al., 2014).

### Encoding models

Linear encoding models were estimated at each lag (-2000 ms to 2000 ms in 25-ms increments) relative to word onset (0 ms) to predict the brain activity for each word from the corresponding contextual embedding. Before fitting the encoding model, we smoothed the signal using a rolling 200-ms window (i.e., for each lag, the model learns to predict the average single +-100 ms around the lag). We estimated and evaluated the encoding models using a 10-fold cross-validation procedure: ridge regression was used to estimate a weight matrix for predicting word-by-word neural signals in 9 out of 10 contiguous training segments of the podcast; for each electrode, we then calculated the Pearson correlation between predicted and actual neural signals for the left-out test segment of the podcast. For each ridge regression model (for each fold, lag, and electrode), the alpha parameter is determined by cross-validation using the “RidgeCV” method from the “himalaya” package (Dupré la Tour et al., 2022). This procedure was performed for all layers of contextual embeddings from each LLM. To control for the different embedding dimensionality across models, we standardized all embeddings to the same size using principal component analysis (PCA) and trained linear encoding models using ordinary least-squares (OLS) regression. The PC features are used by the OLS models only.

### Dimensionality reduction

To control for the different hidden embedding sizes across models, we standardized all embeddings to the same size using principal component analysis (PCA) and trained linear regression encoding models using ordinary least-squares regression, replicating all results (Fig. S2). This procedure effectively focuses our subsequent analysis on the 50 orthogonal dimensions in the embedding space that account for the most variance in the stimulus. We compute PCA separately on the training and testing set to avoid data leakage.

## Data & Code Availability

We have recently made the data publicly available (Zada et al., 2025). We have also provided tutorials for preprocessing the data and training encoding models: https://hassonlab.github.io/podcast-ecog-tutorials. For this specific project, the analysis code is available at https://github.com/hassonlab/247-pickling/tree/scaling-paper-0 and https://github.com/hassonlab/247-encoding/tree/scaling-paper-1.

## Acknowledgments

This work was supported by the National Institutes of Health under award numbers DP1HD091948 (to A.G., Z.H., H.W., Z.Z., B.A., L.N., A.F., and U.H.), R01NS109367 (to A.F.), and R01DC022534 (to S.A.N.), Finding a Cure for Epilepsy and Seizures (FACES), and Schmidt Futures Foundation DataX Fund. Z.H. devised the project, performed experimental design and data analysis, and wrote the article; H.W. devised the project, performed experimental design and data analysis, and wrote the article; Z.Z. devised the project, performed experimental design and data analysis, and critically revised the article; H.G. performed data analysis; B.A. performed data analysis; L.N. performed data analysis; W.D. devised the project; S.D. devised the project; P.D. devised the project; D.F. devised the project; O.D. devised the project; A.F. devised the project; U.H. devised the project, performed experimental design, and critically revised the article; S.A.N. devised the project, performed experimental design, wrote and critically revised the article; A.G. devised the project, performed experimental design, and critically revised the article.

---

**Supplementary Figure 1.**
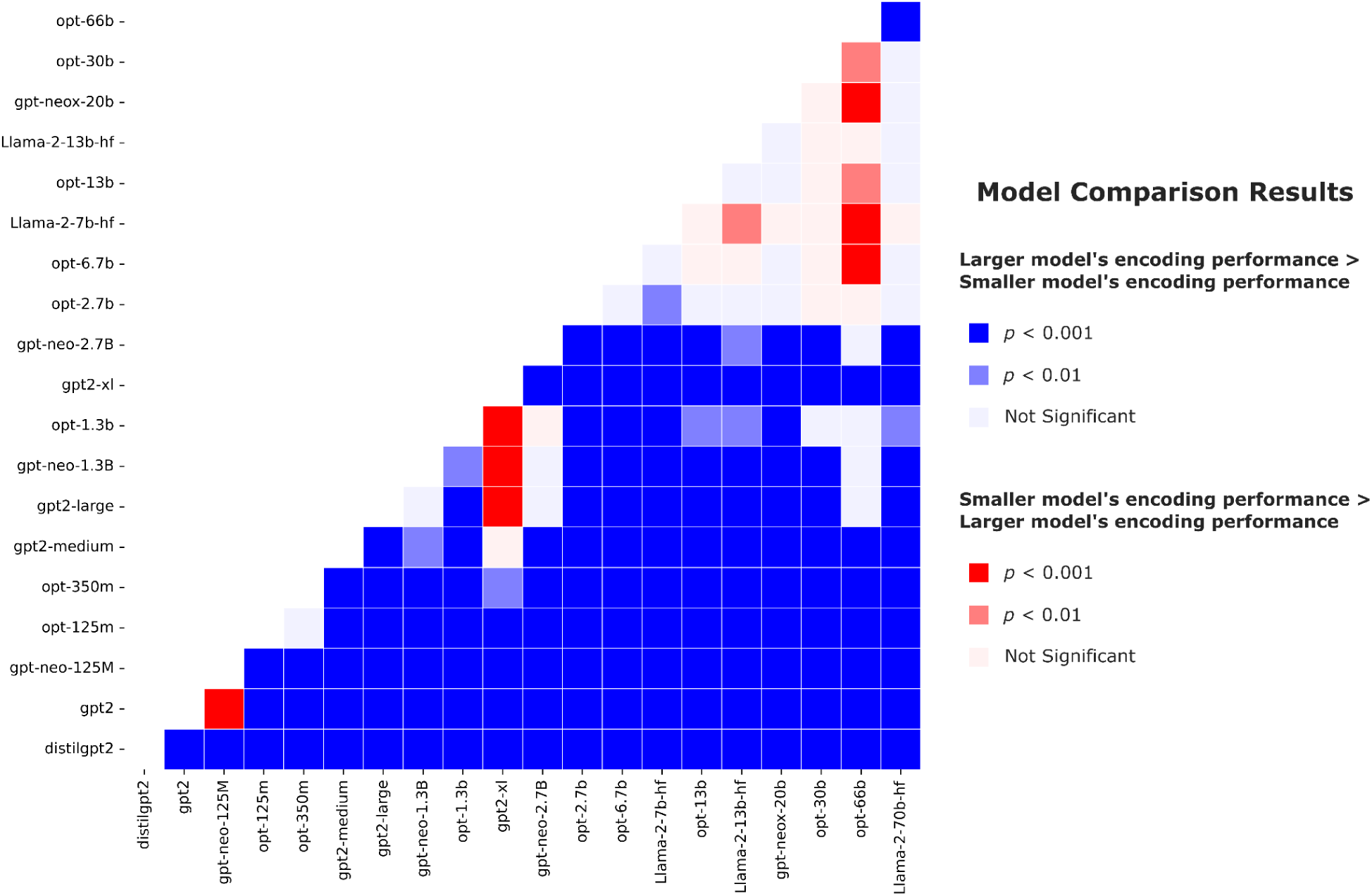
Heatmap comparing encoding performance between models. Encoding performance for a model is represented by the maximum correlation across lags and layers per electrode. Paired t-tests are performed across electrodes (df = 159). The result is blue if t < 0, meaning the larger model (the model with a higher number of parameters, represented on the x-axis) outperforms the smaller model (the model with a smaller number of parameters, represented on the y-axis). The result is red if t > 0, meaning the smaller model outperforms the larger model. The shades of the colors represent significance (p < 0.001 or p < 0.01 or not significant, two-sided, FDR corrected). There is a positive relationship between model size and encoding performance when models are smaller than 3 billion parameters. When models are larger than 7 billion parameters, we observed a plateau in the maximal encoding performance.

**Supplementary Figure 2.**
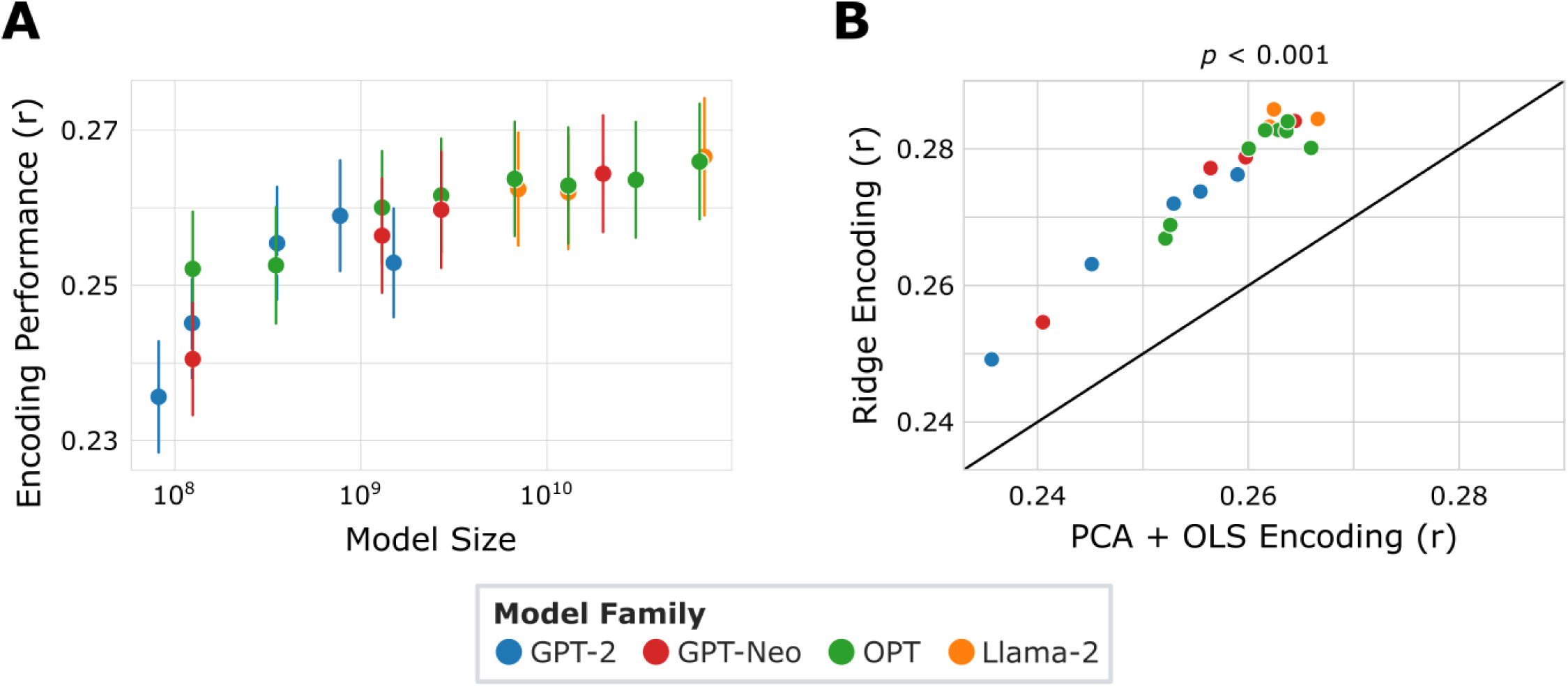
Model performance improves with increasing model size. To control for the different embedding dimensionality across models, we standardized all embeddings to the same size using principal component analysis (PCA) and trained linear encoding models using ordinary least-squares (OLS) regression (cf. Fig. 2). **A.** Replication of Fig. 2B, the relationship between model size (shown on a log scale) and brain encoding performance: encoding performance increases as model size increases. Each data point corresponds to a model. **B.** Ridge regression encoding outperforms PCA + OLS regression encoding for all 20 transformer-based language models (paired two-sided *t*-test across electrodes, df = 159, *p* < 0.001, Bonferroni corrected). Each data point corresponds to a model.

**Supplementary Figure 3.**
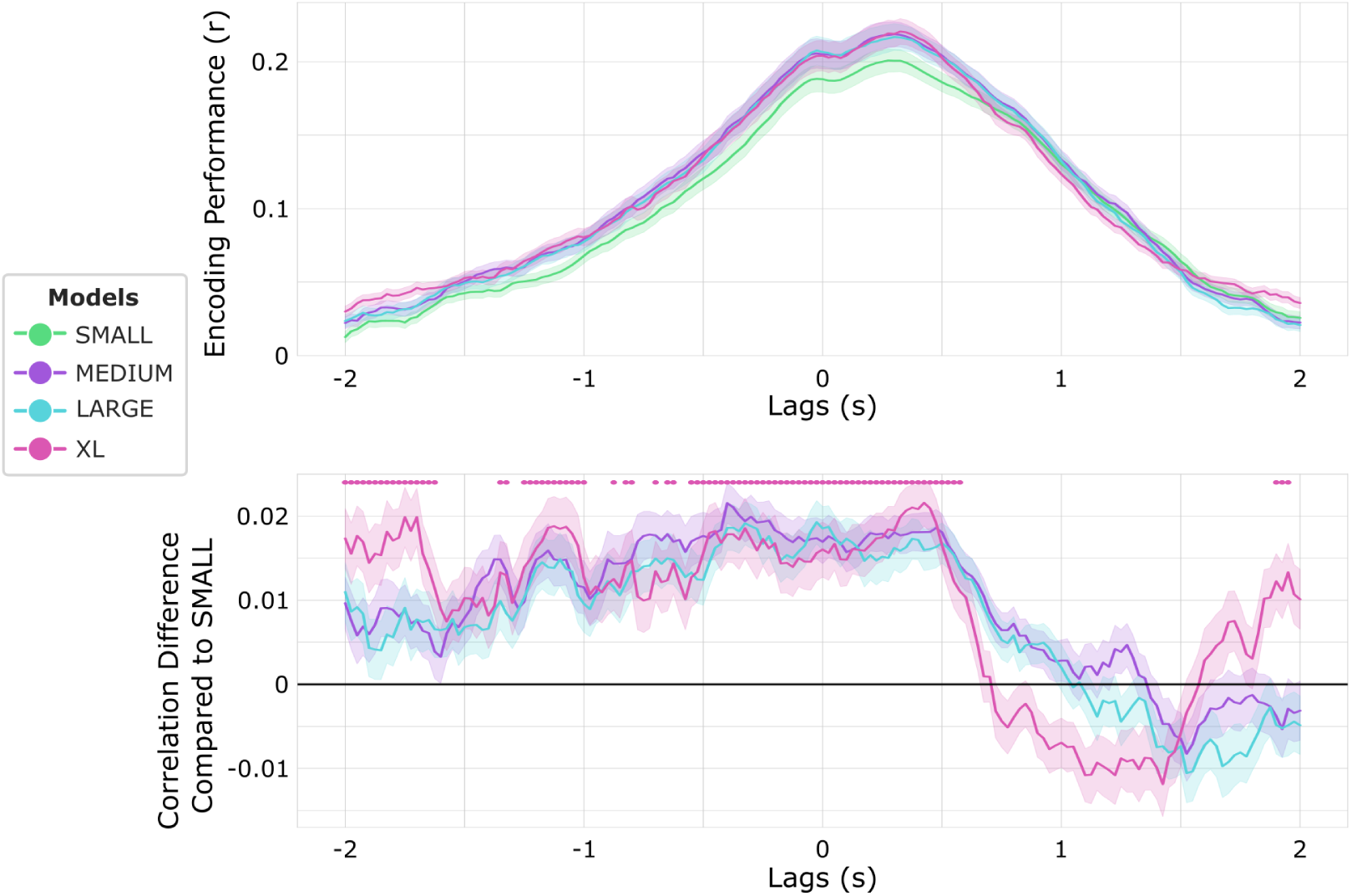
Lag-wise encoding for the GPT-Neo Family. **Top.** Lag-wise encoding for all four models of the GPT-Neo family, averaged across electrodes. The dots represent lags where XL significantly outperformed Small (paired two-sided *t*-test across electrodes, df = 159, *p* < 0.001, Bonferroni corrected). XL significantly outperformed Small in encoding models for most lags from 2000 ms before word onset to 575 ms after word onset. **Bottom.** Lag-wise encoding difference for the three bigger models compared to SMALL, averaged across electrodes.

**Supplementary Figure 4.**
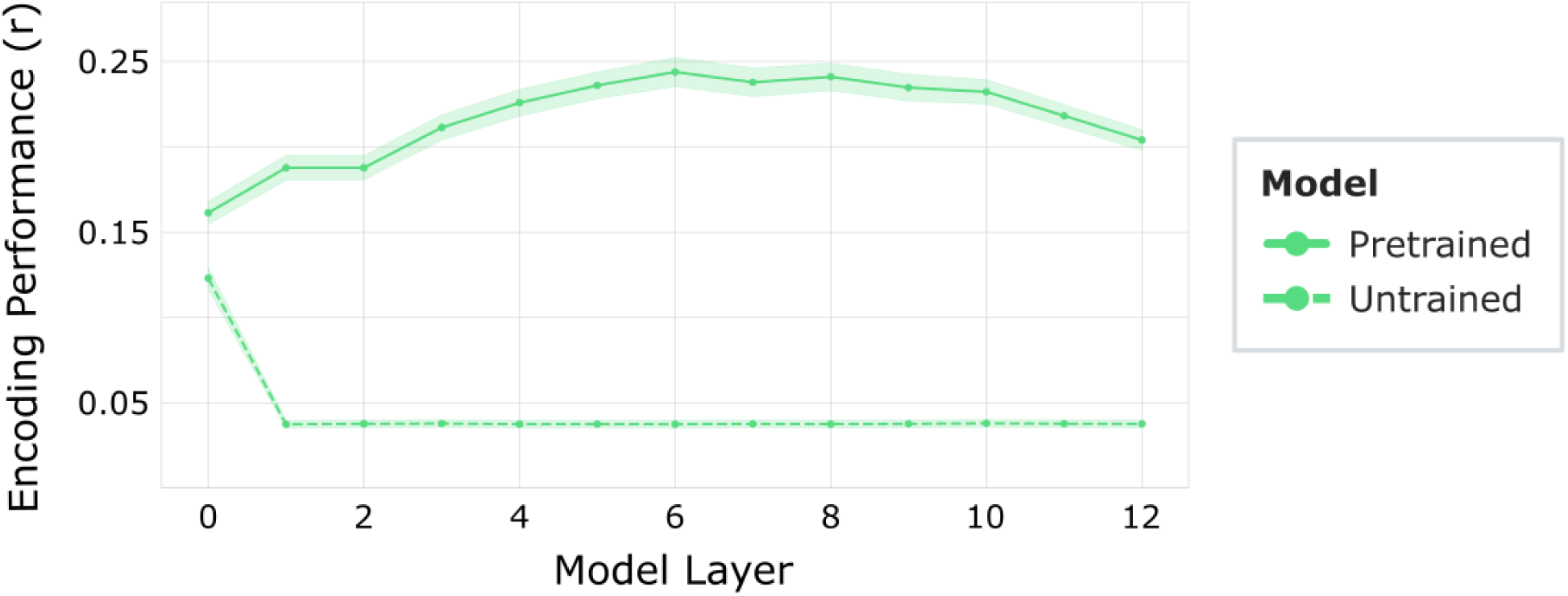
The relationship between encoding performance and layer number for pretrained and untrained SMALL. For pretrained SMALL, encoding performance is best for intermediate layers. For untrained SMALL with randomly initialized weights, encoding performance is best for the 0th layer. Encoding performance is significantly higher for pretrained SMALL than for untrained SMALL for every layer (paired two-sided t-test across electrodes, df = 159, p < 0.001, Bonferroni corrected). The shaded colors represent standard error.

**Supplementary Figure 5.**
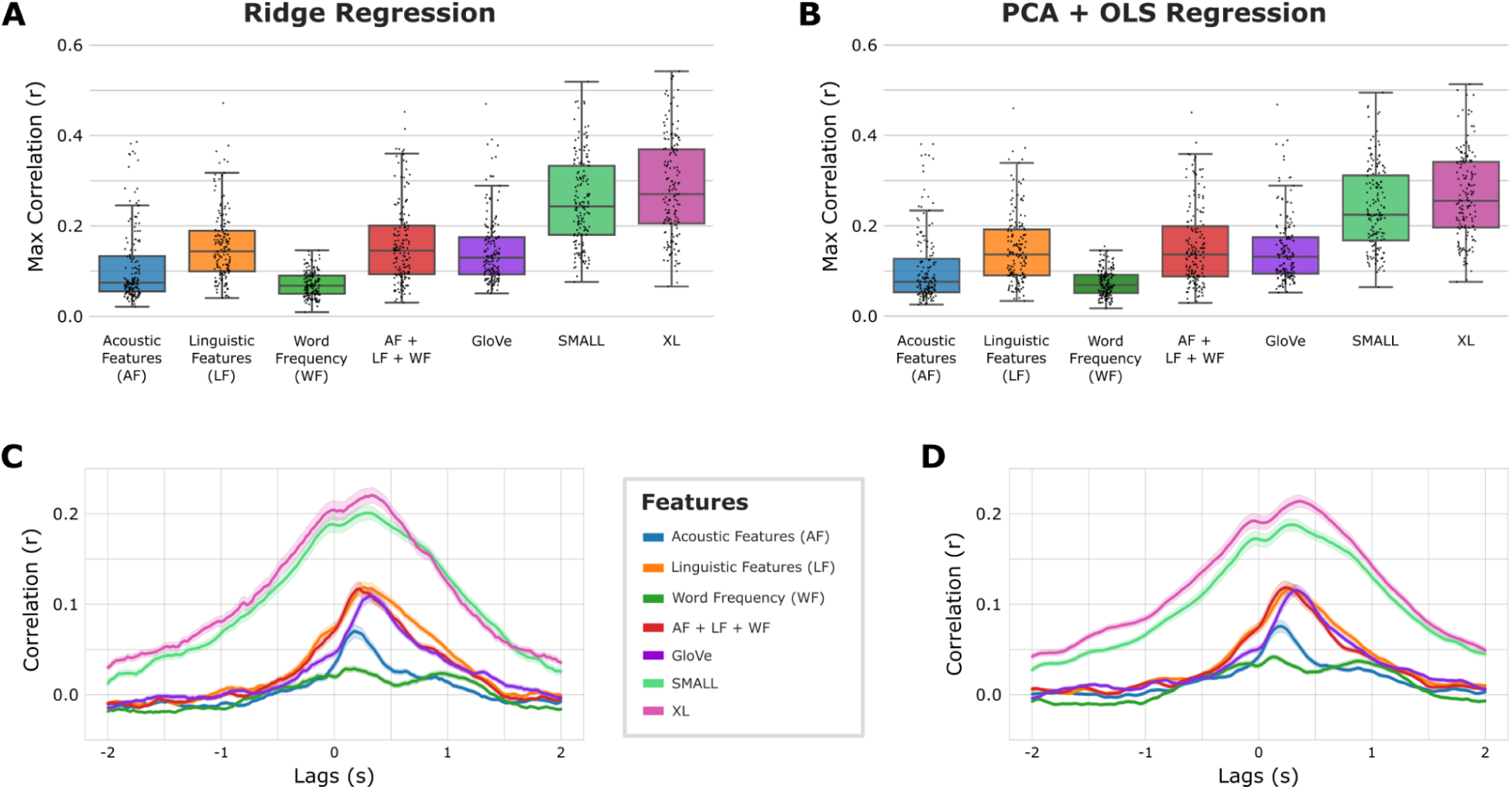
Contextual embeddings from LLMs outperform classical speech features and GloVe embeddings. **A**. Maximum encoding performance (across lags) for acoustic features (60 dimensions), linguistic features (191 dimensions), word frequency (2 dimensions), acoustic features + linguistic features + word frequency (253 dimensions), GloVe (50 dimensions), SMALL (768 dimensions), and XL (6144 dimensions) using ridge regression. SMALL and XL showed significantly better performance than other embeddings (paired two-sided t-test across electrodes, df = 159, p < 0.001). **B**. Replication of A using PCA and OLS regression. To control for the different dimensions of the different feature spaces, we standardized all feature sets to 50 dimensions (2 dimensions for word frequency) using PCA and trained OLS regression encoding models. SMALL and XL showed significantly better performance than other embeddings (paired two-sided t-test across electrodes, df = 159, p < 0.001). **C**. Encoding performance across lags for all features. Shaded colors represent standard error across electrodes. **D.** Replication of C using PCA and OLS regression.

**Supplementary Figure 6.**
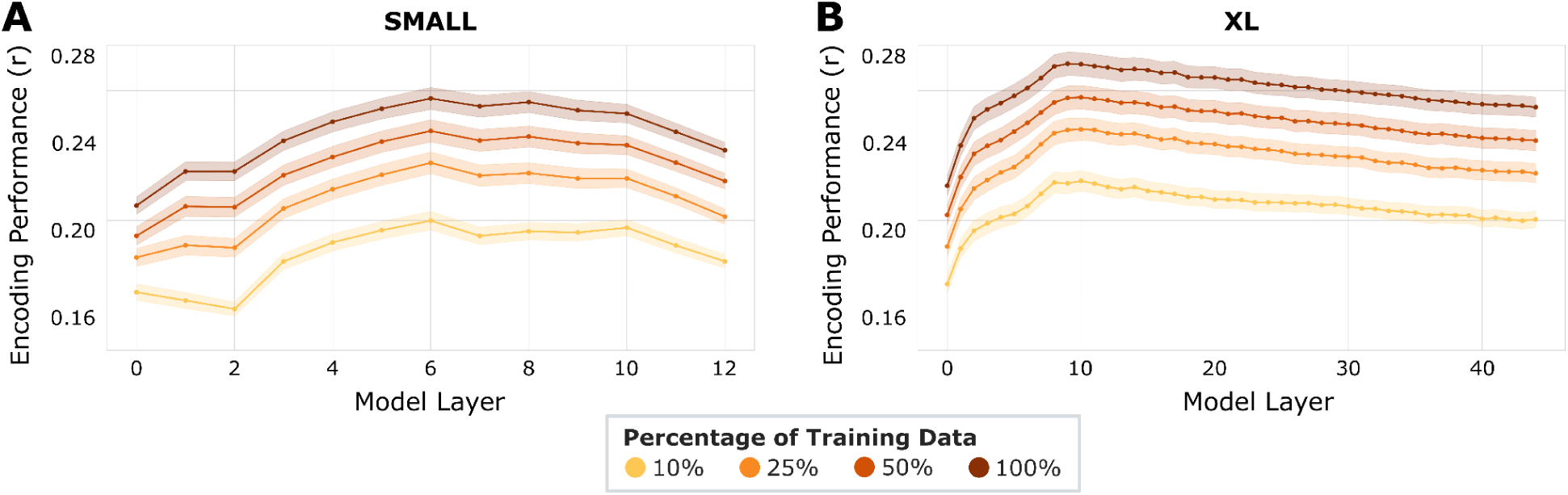
Model performance improves with increasing dataset size. **A.** For SMALL, the relationship between the percentage of training data and brain encoding performance across layers. Encoding performance increases as the training dataset size increases. The shaded colors represent standard error across electrodes. B. Same as A, but for model XL.

**Supplementary Figure 7.**
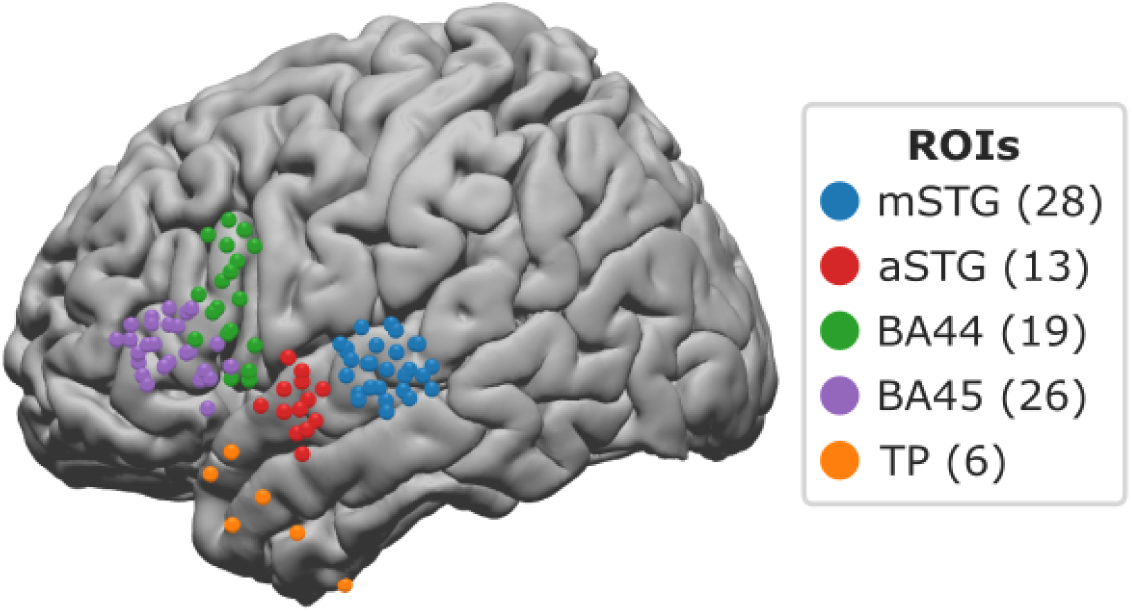
Brain map of electrodes in five regions of interest (ROIs) across the cortical language network: middle superior temporal gyrus (mSTG, n = 28 electrodes), anterior superior temporal gyrus (aSTG, n = 13 electrodes), Brodmann area 44 (BA44, n = 19 electrodes), Brodmann area 45 (BA45, n = 26 electrodes), and temporal pole (TP, n = 6 electrodes).

**Supplementary Figure 8.**
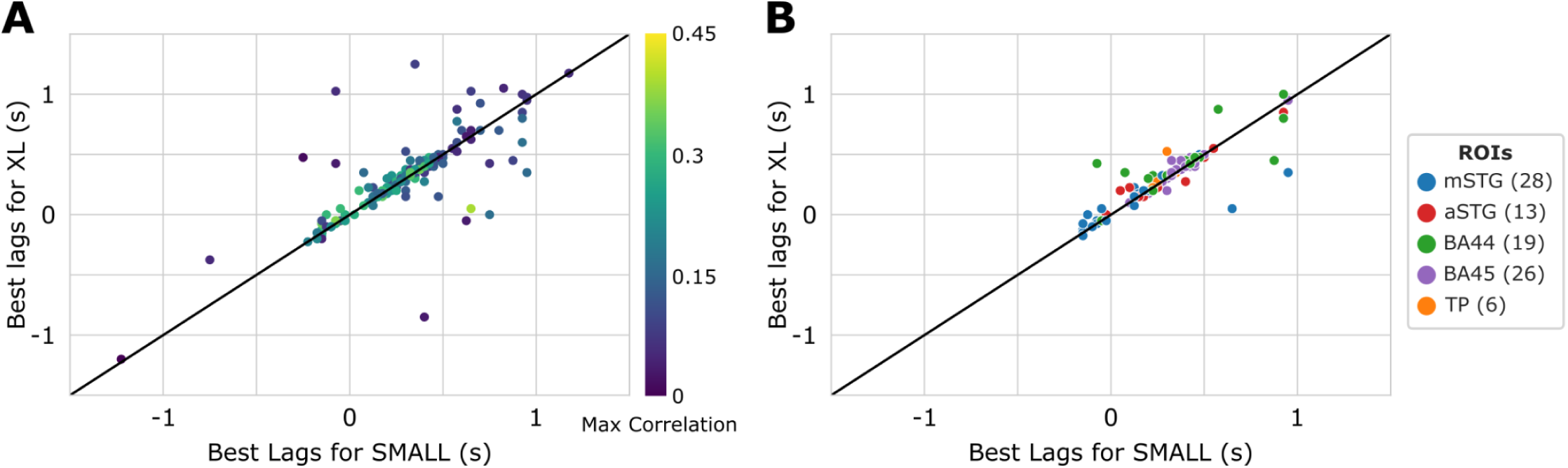
The optimal lags for each electrode do not exhibit significant variation when transitioning between SMALL and XL models. **A.** Scatter plot of best-performing lag for SMALL and XL models, colored by max correlation. Each data point corresponds to an electrode. **B.** Scatter plot of best-performing lag for SMALL and XL models, colored by ROIs. Each data point corresponds to an electrode. Only the electrodes in Fig. S7 are included.

**Supplementary Table 1.**
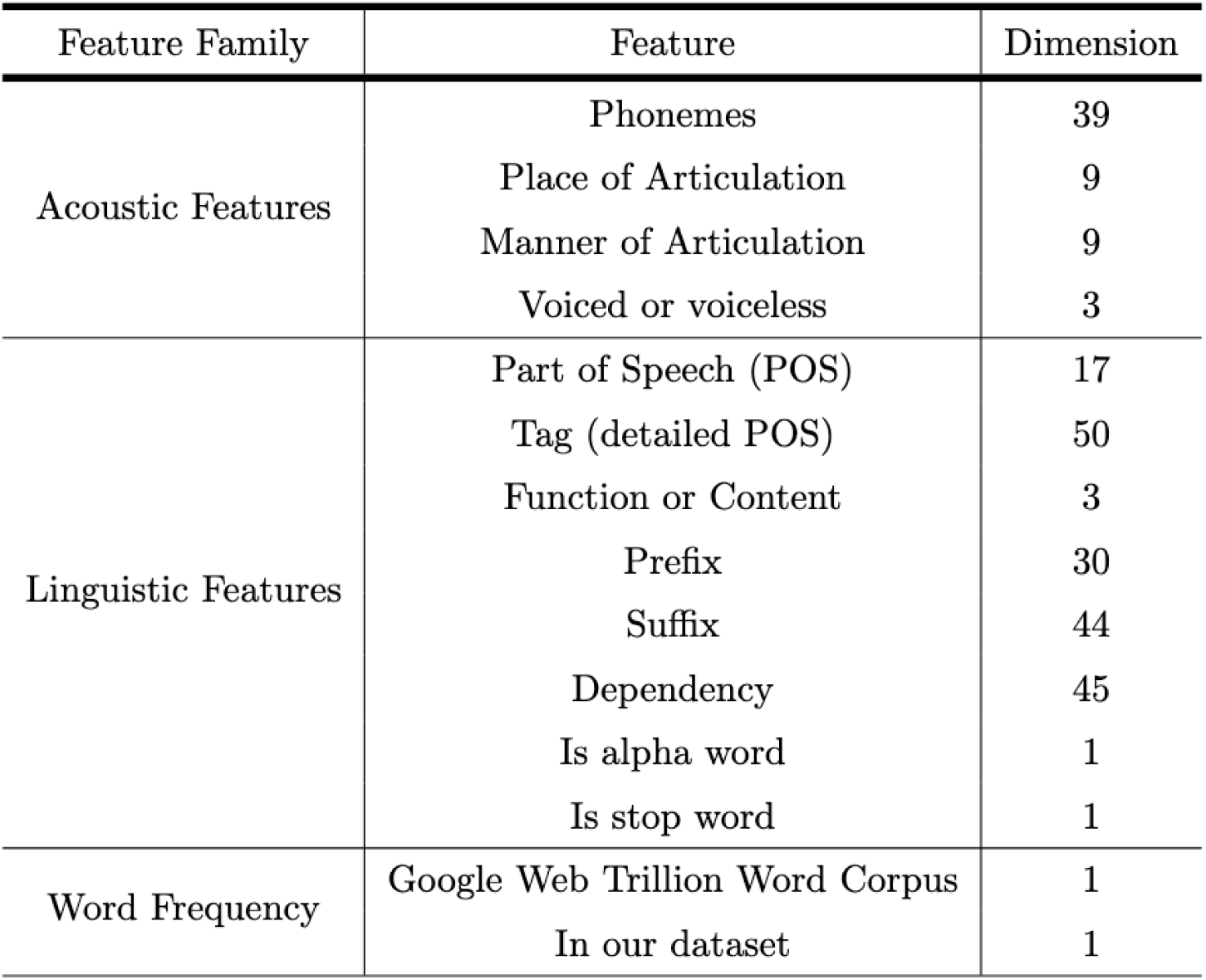
Classic speech features. Lower-level acoustic features include phoneme (39 classes), place of articulation (9 classes), manner of articulation (9 classes), and voiced or voiceless (3 classes). Linguistic features include part of speech (17 classes), tag (50 classes), function or content (3 classes), prefix (30 classes), suffix (44 classes), dependency (45 classes), whether the word is an alpha character (binary), and whether the word is a stop word (binary). Word frequency includes the frequency of the word in the Google Web Trillion Word Corpus and in our dataset.

| Feature Family | Feature | Dimension |
| --- | --- | --- |
| Acoustic Features | Phonemes | 39 |
|  | Place of Articulation | 9 |
|  | Manner of Articulation | 9 |
|  | Voiced or voiceless | 3 |
| Linguistic Features | Part of Speech (POS) | 17 |
|  | Tag (detailed POS) | 50 |
|  | Function or Content | 3 |
|  | Prefix | 30 |
|  | Suffix | 44 |
|  | Dependency | 45 |
|  | Is alpha word | 1 |
|  | Is stop word | 1 |
| Word Frequency | Google Web Trillion Word Corpus | 1 |
|  | In our dataset | 1 |

**Supplementary Table 2.** Summary statistics and paired *t*-test results for maximum correlations between SMALL and XL models across five regions of interest. Encoding performance for the XL model significantly surpassed that of the SMALL model in whole brain, mSTG, aSTG, BA44, and BA45.

| ROI | electrode num | SMALL corr mean | SMALL corr std | XL corr mean | XL corr std | corr diff *t*-value | corr diff *p*-value |
| --- | --- | --- | --- | --- | --- | --- | --- |
| all | 160 | 0.255 | 0.097 | 0.284 | 0.109 | 16.791 | **4.770e<sup>-37</sup>** |
| mSTG | 28 | 0.335 | 0.087 | 0.377 | 0.098 | 11.584 | **5.538e<sup>-12</sup>** |
| aSTG | 13 | 0.274 | 0.096 | 0.311 | 0.108 | 6.111 | **5.247e<sup>-5</sup>** |
| BA44 | 19 | 0.224 | 0.068 | 0.257 | 0.082 | 6.340 | **5.663e<sup>-6</sup>** |
| BA45 | 26 | 0.289 | 0.105 | 0.316 | 0.113 | 6.107 | **2.205e<sup>-6</sup>** |
| TP | 6 | 0.243 | 0.086 | 0.263 | 0.086 | 1.991 | 0.103 |

**Supplementary Table 3.**
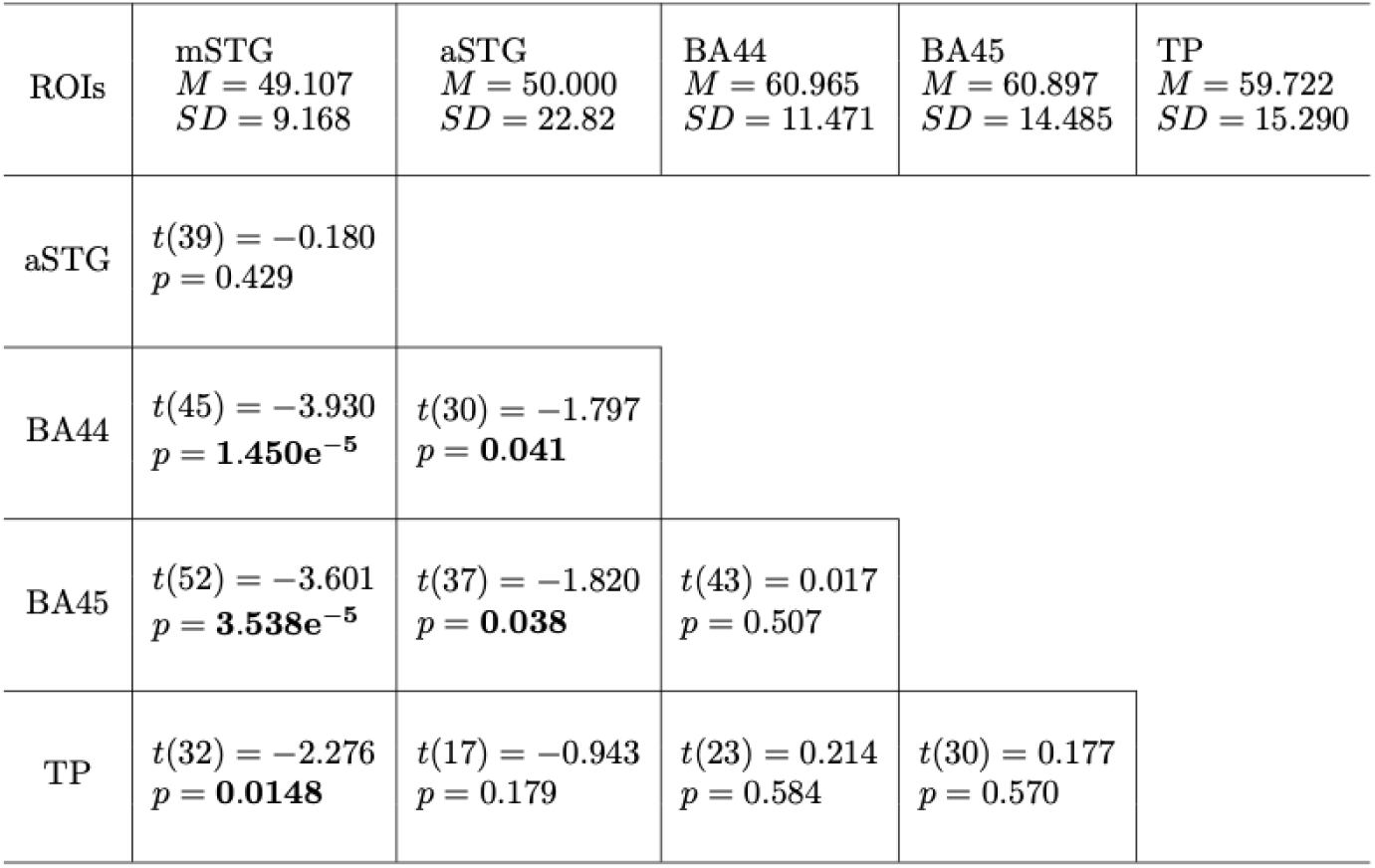
Summary statistics and paired *t*-test results for best-performing layers (in percentage) for the SMALL model across five regions of interest. The best-performing layer (in percentage) occurred earlier for electrodes in mSTG and aSTG and later for electrodes in BA44, BA45, and TP.

